# New Insights Into The Melanin Structure Of *Lomentospora prolificans*

**DOI:** 10.1101/2024.11.01.621558

**Authors:** Livia C. Liporagi-Lopes, Christine Chrissian, Arlind Kacirani, Emma Camacho, Ruth E. Stark, Arturo Casadevall

## Abstract

*Lomentospora prolificans* is a filamentous fungus with a global distribution, manifesting particularly higher prevalence in human-impacted environments. This organism is associated with a wide spectrum of human infections, especially in immunosuppressed individuals, for whom it causes severe and debilitating illnesses with high morbidity and mortality that are compounded by its pan-resistant profile with respect to antifungal drugs. Melanin is a ubiquitous pigment among fungi with a broad range of actions that include promoting fungal virulence. Although melanin is one of the most studied virulence factors in pathogenic fungi, relatively little is known about the chemistry of this pigment in *L. prolificans.* In the current study we characterized *L. prolificans*-associated melanin using chemical, biological, biophysical and structural techniques, also assessing the impact of inhibitors of distinct melanization pathways. Our results reveal that this pathogenic fungus makes multiple types of melanin pigments and suggests the possibility of a new type of melanin, which is synthesized together with a mixture of DHN-, DOPA- and pyomelanin types. These insights enhance our understanding of *L. prolificans’* virulence mechanisms, paving the way for potential therapeutic interventions.

## Introduction

*Lomentospora* (formerly *Scedosporium*) *prolificans* is a fungal pathogen that causes severe disease in immunocompromised patients, with most cases occurring in persons with malignancy or solid organ transplantation (1). *L. prolificans* isolates are pan-resistant to all systemically active antifungals, including azoles, terbinafine and amphotericin B, leading to high mortality rates in these patients (1–3).

Melanins are natural biopolymers that serve different biological functions, ranging from electromagnetic energy capture to protection of microorganisms from adverse environmental conditions; they contribute to the virulence of many pathogenic fungi and bacteria (4,5). Despite their biological relevance, the fundamental structures of these pigments remain unsolved because they are insoluble amorphous solids that cannot be crystallized. Melanins are composed of covalently linked phenol- and indole-type compounds; fungal melanins are closely associated with lipid, carbohydrate, and protein constituents (6). These pigments display heterogeneity in chemical composition, size, color, and function (4).

There are several classes of melanins, each produced through different biosynthetic pathways: 1) eumelanin, a black–brown pigment that is produced by the polymerization of catecholamine oxidation products, including reactive quinone and indole intermediates; 2) pheomelanin, a red–yellow pigment produced by the polymerization of oxidized catecholamines with cysteine-containing amino acids; 3) pyomelanin, a light-brown pigment derived from tyrosine catabolism and the polymerization of homogentisic acid; and 4) dihydroxynaphthalene melanin (DHN-melanin), a black-brown pigment produced from polymerized acetyl-CoA via the polyketide synthesis pathway. The predominant melanins found in the fungal kingdom are DOPA (7) and DHN melanins (7). In addition, there is evidence that some fungi produce pyomelanins that result from the breakdown of aromatic amino acids, particularly tyrosine (7). Some fungal species synthesize more than one type of melanin.

For example, *Fonsecaea pedrosoi* (7), *Colletotrichum lagenarium* (8), and *Magnaporthe grisea* (9) each synthesize DHN-melanin. DOPA-melanin is produced by species such as *Aspergillus flavus* (10), *Aspergillus nidulans* (11), and *Cryptococcus gattii* (12). Other species can produce more than one type of melanin, including *Aspergillus fumigatus* (DOPA-, DHN- and pyomelanin), *Cryptococcus neoformans* (DOPA and pyomelanin (7), *Aspergillus niger* (DOPA- and DHN-melanin) (7), *Exophiala (Wangiella) dermatitidis* (DOPA- and DHN-melanin), *Alternaria alternata* (13) and *Histoplasma capsulatum* (DOPA-, DHN- and pyomelanin) (7).

Whereas *L. prolificans* conidia have been reported to synthesize DHN-melanin (14), the structure and synthesis of the melanin pigment in this fungal species remains inadequately studied. The few studies carried out thus far do not provide comprehensive information about the structure, synthesis and function of this pigment in *L. prolificans* (14–16). Furthermore, analyses conducted to study melanin in *L. prolificans* have relied on indirect experiments and tools, such as using monoclonal antibodies against melanin-biosynthetic enzyme tetrahydroxynaphthalene reductase (16), or generating enzyme-deficient mutants with abnormal pigmentation due to disruption in normal DHN-melanin biosynthesis by a random mutation (UV mutagenesis) (15). Additionally, elemental analysis alone is not sufficient to differentiate between pigment types such as DHN- and eumelanin. Given these circumstances, a study of melanogenesis inhibition using inhibitors specific to each biosynthetic pathway has the potential to yield more informative results. In the current report, we revisit the problem of melanization in *L. prolificans* and present evidence that the organism produces a complex array of melanin pigments.

## Materials and Methods

### Fungal strains, cell growth and culture conditions

The *Lomentospora prolificans* strain used in this study is a naturally antifungal-resistant clinical isolate from an 11-year-old boy in Australia (23) (*L. prolificans* AMMRL 140.04, catalogue no. 90853; ATCC, Manassas, VA). AMMRL 140.04 was grown on potato dextrose (PD) broth (rich medium) in flasks for 14 d at 30°C to collect sufficient fungal biomass for the experiments. To obtain conidia, cells were grown on potato dextrose agar (PDA) for 7 d at 30°C prior to harvesting by scraping the plate surface with phosphate buffered saline (PBS).

### Reagents

The following melanin inhibitors were used: DHN-melanin inhibitors pyroquilon (Sigma-Aldrich # 45650) and tricyclazole (Sigma-Aldrich, # 45808) (13,17); the DOPA-melanin inhibitor niacin (nicotinic acid, Sigma-Aldrich, # N0761) (18); and the pyomelanin inhibitor nitisinone (Sigma-Aldrich, # SML0269) (19). Other reagents from Sigma-Aldrich included ABTS (2,2′-Azino-bis(3-ethylbenzothiazoline-6-sulfonic acid) diammonium salt (#10102946001) and homogentisic acid (# 678626) (20), each used as melanin substrate precursors in particular experiments.

### Melanin Ghost Isolation

The following protocol was used as previously described (1996) (21). Melanized *L. prolificans* cells were collected by centrifugation at 1,000 *g,* washed with phosphate-buffered saline (PBS), suspended in a 1 M sorbitol/0.1 M sodium citrate pH 5.5 solution, and incubated at 30 °C for 24 h with 10 mg/mL of lysing enzymes from *Trichoderma harzianum* (Sigma-Aldrich). The enzyme-digested cells were collected again by centrifugation at 10,000 *g* for 10 min and washed repeatedly with PBS until the supernatant was nearly clear. Proteinaceous materials were denatured by incubating the pellets with 4 M guanidine thiocyanate in a rocker for 12 h at room temperature. The recovered debris was then collected by centrifugation, washed 2-3 times with ∼20 mL of PBS, and incubated for 4 h at 65 °C in 10 mL of buffer (10 mM pH 8.0 Tris-HCl, 5 mM CaCl2, 5% sodium dodecyl sulfate) containing 1 mg/mL of proteinase K (Boehringer, Mannheim, Germany). After washing three times with ∼20 mL of PBS, lipids were extracted three successive times by the Folch method using a chloroform, methanol, and saline mixture at a ratio of 8:4:3 (v/v/v). The remaining material was suspended in 20 mL of 6 M HCl and boiled for 1 h to hydrolyze remaining cellular contaminants associated with the melanin. The resulting acid-resistant insoluble material was dialyzed against distilled water for 14 d with daily water changes and lyophilized. Elemental analysis was performed to determine the percentage of carbon, nitrogen, oxygen, and hydrogen in the melanin pigments (Intertek, Whitehouse, NJ) (22).

### Measurement of fungal growth and pigment production

The capacity of *L. prolificans* to grow and produce pigment in the presence of melanin inhibitors was evaluated. To monitor the growth capacity of the strains, conidia (1 x 10^5^ per condition) were inoculated to media, incubated at 30°C with shaking at 180 rpm, and then monitored over time by measuring the absorbance at 600 nm (23). Fungal cell viability was determined by the XTT reduction assay, measuring the optical absorbance at 490 nm (24,25). To analyze melanin production, 1 x 10^7^ conidia/ml were suspended in aqueous solution and analyzed by measuring their absorbance over a wavelength range of 400-700 nm under different growth conditions. Additionally, suspensions of 1 x 10^7^ conidia/ml from different cultivation times were solubilized in 1 M NaOH/10% DMSO for 2 h at 80°C, then centrifuged at 1,200 g for 10 min at room temperature. The supernatants were transferred to fresh tubes, filtered through a Millipore filter (pore size, 0.45 μm), and analyzed by measuring the absorbance at 405 nm (26,27).

### Extracellular phenol oxidase production (Laccase Activity)

Laccase activity was determined by evaluating the oxidation of ABTS (28), a nonphenolic dye that is oxidized by laccase into a more stable cation radical. The concentration of the cation radical responsible for the intense blue-green color then provides a quantitative measure of enzyme activity [13]. The assay mixture contained 0.5 mM ABTS, 0.1 M sodium acetate (pH 4.5), and 0.1 ml of enzyme. The activity was measured after 7 d of fungal growth.

### Phagocytic and killing assays

Primary macrophages were obtained from C57BL/6 mouse bone marrow progenitors. Mice were euthanized, the skin of each leg was peeled back, and muscle was cut away to reveal the bone. An L929 cell-conditioned supernatant medium was obtained by collecting supernatants from L929 cells at confluence. Once the skin and muscle were removed, the tibia and femur were removed and placed in Dulbecco modified Eagle feeding medium (DMEM) (20% L929 cell supernatant, 10% fetal bovine serum, 0.1% β-mercaptoethanol, 1% penicillin-streptomycin, 1% minimal essential medium nonessential amino acids, 1% GlutaMAX, 1% HEPES buffer). The marrow was then flushed out with feeding medium, filtered through 100 μm pore filters to remove aggregates or bone debris, and centrifuged at 350 *g* for 5 min at 4°C. The supernatant was then discarded, and the pellet was suspended in 120 ml of feeding medium. The resuspended solution was seeded onto 100 mm tissue culture-treated petri dishes (Corning) and incubated at 37°C for 6 d (29). Mammalian bone marrow-derived macrophage cells (BMDM) were seeded onto the wells of a 24-well plate containing BMDM feeding medium at a density of 2 × 10^5^/well. The BMDM were incubated overnight at 37°C with gamma interferon (IFN-γ; Bioscience) and lipopolysaccharide (LPS; Bioscience) at 100 U/ml and 0.5 μg/ml, respectively. BMDM were infected at a multiplicity of infection of 3:1 (conidia cells/BMDM). BMDM were incubated for 2 h at 37°C before imaging (29). For the killing assay, after the interaction time, the wells were first washed with PBS (to remove any extracellular or adherent conidia), and macrophages were lysed by adding cold sterile deionized water. Aliquots were plated on PDA plates, and the plates were incubated at 30°C for 5 d.

### Zeta potential

The charge of each sample was measured using a Zeta potential analyzer (ZetaPlus, Brookhaven Instruments Corp., Holtsville, NY), as described previously (30). Measurements were done at 25°C and expressed in mV units.

### Transmission electron microscopy (TEM)

*L. prolificans* melanin particles were fixed in 2.5% (v/v) glutaraldehyde in 0.1 M sodium phosphate buffer (pH 7.3) for 24 h at 4°C. Samples were encapsulated in 3% (w/v) low-melting-point agarose prior to being processed in Spurr resin following a 24-h schedule on a Lynx tissue processor (secondary 1% osmium tetroxide fixation, 1% uranyl acetate contrasting, ethanol dehydration, and infiltration with acetone/Spurr resin). Additional infiltration was conducted under vacuum at 60°C before the samples were embedded in TAAB capsules (TAAB Laboratories, Amersham Biosciences, UK) and polymerized at 60°C for 48 h. Semithin survey sections (0.5 μm thick) were stained with 1% toluidine blue to identify the areas with the best cell density. Ultrathin sections (60–90 nm) were prepared with a Diatome diamond knife on a Leica UC6 ultramicrotome and stained with 2% uranyl acetate in 50% methanol, followed by lead citrate for examination with a JEOL 1200EX TEM (Analytical Imaging Facility at The Albert Einstein College of Medicine) or a Philips CM120 TEM operating at 80 kV (Microscopy Facility at the Johns Hopkins School of Medicine). Negative staining of the fractions from OptiPrep density gradients was performed by adsorbing 10 μl of each fraction to glow-discharged 400 mesh ultra-thin carbon-coated grids (EMS CF400-CU-UL) for 2 min, followed by three quick rinses with Tris-buffered saline solution and staining with 1% uranyl acetate containing 0.05% Tylose. Images were captured with a Gatan Orlus Digital 2K × 2K CCD camera or an AMT XR80 high-resolution (16-bit) 8-megapixel camera.

### Electron Paramagnetic Resonance (EPR)

Melanin particles (∼3 mg) were suspended in 150 µl distilled water, sonicated for 1 min using a horned sonicator, and examined by EPR spectroscopy as described by Camacho *et al*., 2019 (31). EPR spectra were recorded at the Instrumentation Facility of the Department of Chemistry, Johns Hopkins University, Baltimore, USA, using a Bruker EMX EPR spectrometer operating at 77 K in X-band mode.

### Fourier Transform Infrared (FTIR) spectroscopy

Molecular groups in the melanins (∼5 mg) from *L. prolificans* were characterized by FTIR using a Nicolet Nexus 670 spectrometer coupled with a Smart Gold Gate KRS-5 accessory and OMNIC software (ThermoFisher Scientific, USA). FTIR spectra were recorded between 4000 and 400 cm^-1^ in transmittance mode by averaging 32 scans. Spectra resolution was 4 cm^-1^. A standard DOPA-melanin spectrum was also obtained using *Cryptococcus neoformans* melanin.

### Solid-state NMR (ssNMR)

ssNMR experiments were carried out on a 600 MHz Varian (Agilent) DirectDrive2 (DD2) spectrometer using a 1.6-mm FastMAS probe (Agilent Technologies, Santa Clara, CA) with a magic-angle spinning (MAS) rate of 15.00 ± 0.02 kHz and a nominal spectrometer-set temperature of 25°C. Average post-lyophilization sample masses were ∼5 mg for the melanin ghosts. Typical 90° pulse lengths for the ^13^C Cross Polarization Magic-Angle Spinning (CPMAS) NMR experiments were 1.2 μs for ^1^H and 1.4 μs for ^13^C, respectively; a ^1^H-^13^C mixing time 1 ms was used. As described previously (18), ^1^H decoupling was applied during signal acquisition using the small-phase incremental alternation pulse sequence with a field strength of 104 kHz. Recycle delays of 3 s were inserted between 4096 successive acquisitions for each melanin ghost ^13^C NMR spectrum.

### Statistical Analysis

Statistical analyses were done using GraphPad Prism version 8.00 for Mac OSX (GraphPad Software, San Diego, CA, USA). One-way analysis of variance with a Kruskal-Wallis nonparametric test was used to compare the differences between groups, and individual comparisons of groups were performed using a Bonferroni post-test. The Student’s *t* test was used to compare the number of colony forming units across different groups. The 90–95% confidence interval was determined in all experiments.

## Results

### Biological and Structural Characterization of *L. prolificans* Melanins

#### Impact of Melanin Inhibitors on Cell Viability and Pigmentation

To determine whether the melanin inhibitors used to inhibit melanization impacted the viability of the fungal cells, *L. prolificans* growth curves were generated in the presence of different concentrations by measuring absorbance at 600 nm. Some melanin inhibitors, such as pyroquilon and nitisinone, reduced *L. prolificans* growth/viability significantly at doses higher than 30 µg/ml and in a dose-dependent manner (**Figure 1A**). Based on this finding, we designated 30 µg/ml as the concentration acting directly on the melanin synthesis pathways rather than affecting other biological processes that could alter melanin synthesis indirectly as a secondary effect.

**Figure 1.**
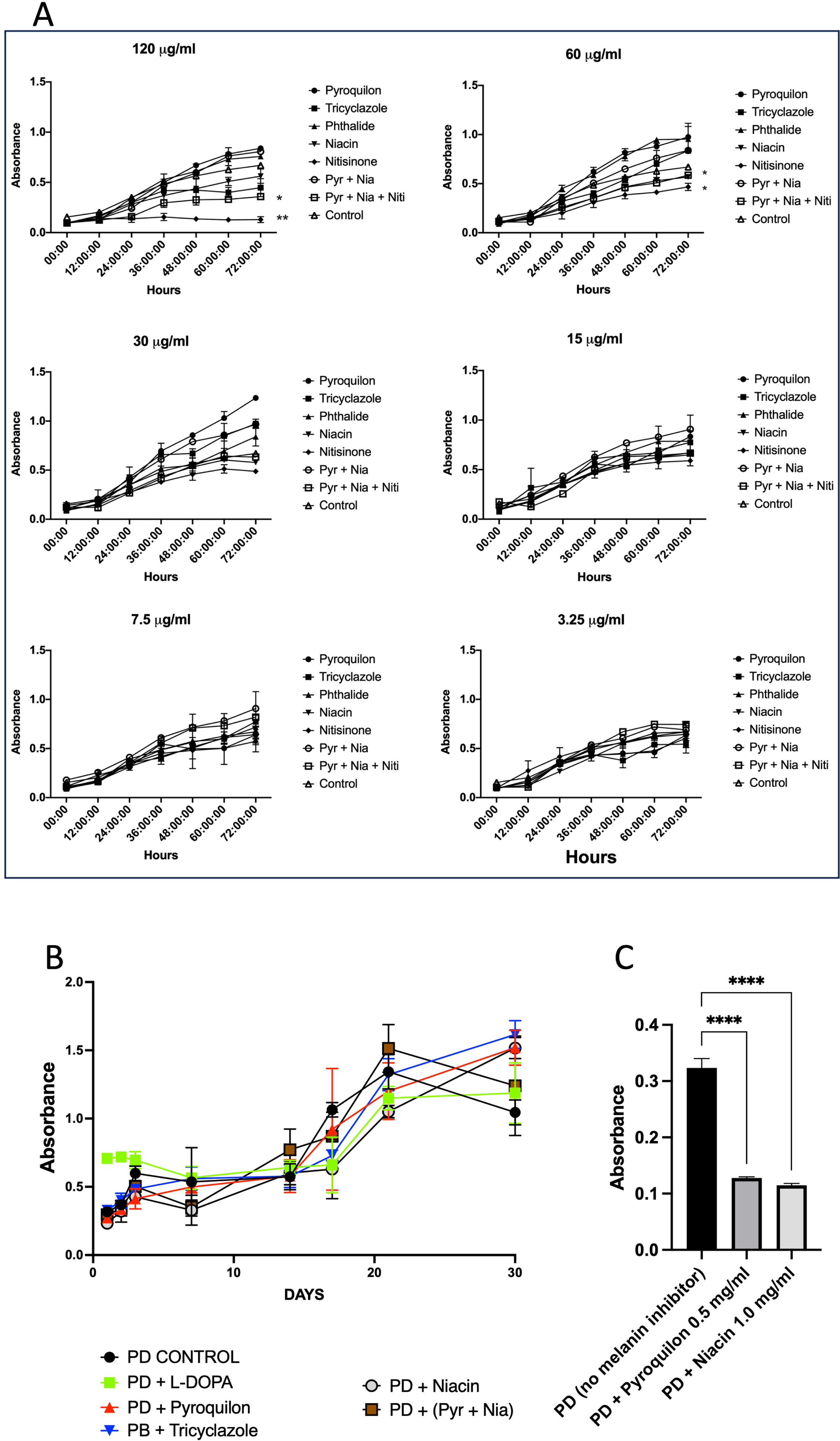
*L. prolificans* growth and absorption curves for different concentrations of melanization inhibitors. Different concentrations of melanin inhibitors were used to evaluate the growth of *L. prolificans* conidia for 3 days, measured by absorbance at 600 nm (A). Standard deviations are calculated from three independent experiments. *L. prolificans* cultures were grown for 30 days at 30°C without or with 30 µg/ml of the melanin inhibitors. The cell viability was measured by XTT at 450 nm (B). The experiments were repeated three times in duplicate. Standard deviations are calculated from three independent experiments. High doses of some melanin inhibitors (0.5 and 1.0 mg of each) were tested in PD liquid media (B), and absorbance was measured at 405 nm. This experiment was repeated three times in duplicate; standard deviations were calculated from three independent experiments. Pyr + Nia: Pyroquilon + Niacin; Pyr + Nia + Niti: Pyroquilon + Niacin + Nitisinone. * *p* < 0.05, ** *p* < 0.005, **** *p* < 0.0001.

Growth of *L. prolificans* in the presence of the various inhibitors (30 µg/ml) was measured for longer intervals of time by XTT reduction (24). The growth curves demonstrated that none of the melanin inhibitors significantly altered the growth/viability of *L. prolificans* at the concentrations tested after 30 d of incubation (**Figure 1B**).

To determine if higher doses of melanin inhibitors affected the fungal pigmentation, *L. prolificans* cells were grown with 0.5 and 1 mg/ml of the melanin inhibitors pyroquilon (DHN-melanin inhibitor) and niacin (DOPA-melanin inhibitor) in PD medium (**Figure 1C**). Under these conditions, the cells manifested a significant decrease in their pigmentation and viability (data not shown). There was no growth observed with 1.0 mg/ml of pyroquilon.

*L. prolificans* cultures grown in different media displayed varying colors (**Figure 2A**). After 14 d of growth on PD medium, both the cells and the supernatant were dark. Addition of 30 µg/ml of different melanin inhibitors changed the color of the fungal cells to a brownish coloration with pyroquilon and tricyclazole addition (both DHN-melanin inhibitors), as well as for the combined treatment of pyroquilon and niacin (DHN- and DOPA-melanin inhibitors, respectively) (**Figure 2A**). Whereas addition of 30 µg/ml of different melanin inhibitors changed the color of the *L. prolificans* cells, none abolished coloration (**Figure 2A**).

**Figure 2.**
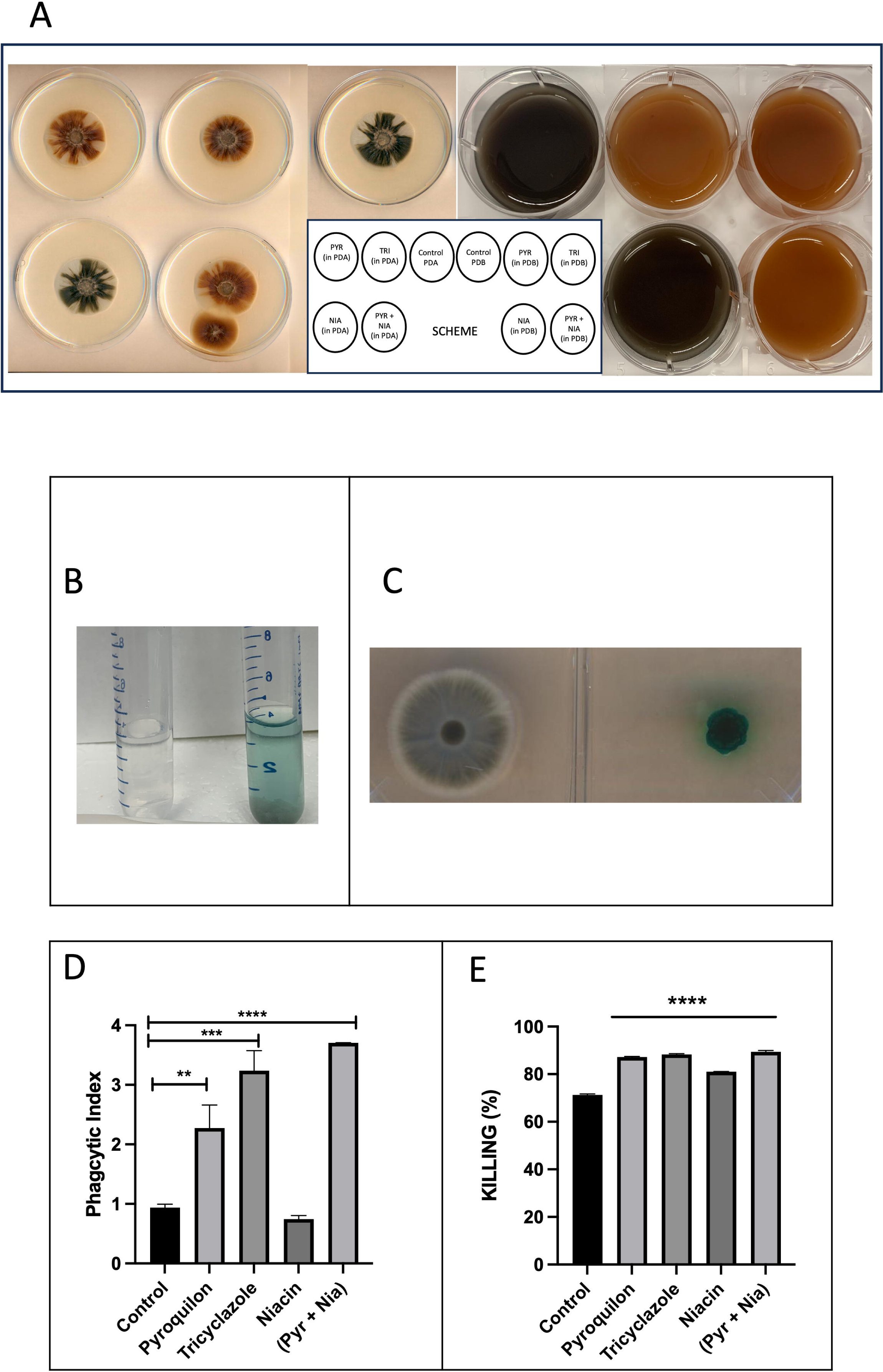
Biological findings related to melanin production by *L. prolificans* cells. *L. prolificans* cultures grown for 14 days at 30°C in the absence or presence of 30 µg/ml of one or more melanin inhibitors in PD agar and broth, respectively, showed different pigment colors **(A).** The efficiency of extracellular phenol oxidase production was determined using the phenol oxidase assay and ABTS as substrate, observing the development of a colored zone in the supernatant (broth media) **(B)** or around fungal colonies (agar media) **(C).** *L. prolificans* conidia grown for 7 days at 30°C in the absence or presence of 30 µg/ml melanin inhibitors were incubated for 2 hours with BMDM cells in DMEM at 37°C/10% CO2, allowing for analysis of phagocytosis **(D)** and killing **(E)** functions. Representative data and standard deviations from three independent experiments are shown. Pyroquilon (PYR), tricyclazole (TRI), phthalide (PHT), niacin (NIA) and pyroquilon + niacin (PYR + NIA). ** *p* < 0.005, *** *p* < 0.0005, **** *p* < 0.0001.

#### Phenol Oxidase activity

Although *L. prolificans* melanin was reported as a DHN-melanin, genes encoding for laccase are found in the *L. prolificans* genome (maker-Scaffold_11-augustus-gene-0.4679-mRNA-1,maker-Scaffold_3-snap-gene-0.2128-mRNA-1 and maker-Scaffold_703-augustus-gene-0.4874-mRNA-1). DOPA-melanin is synthesized by laccases and its production requires exogenously supplied phenolic precursors. Therefore, *L. prolificans* culture supernatants were tested for extracellular phenol oxidase activity using ABTS on both PD and PDA culture media (**Figure 2B**), showing activity by the development of colored zone (blue) in the supernatant or around fungal colonies, a positive indicator for phenol oxidase activity (**Figure 2C**).

#### Phagocytosis and killing assays

To evaluate the interaction between *L. prolificans* cells and BMDM, fungal conidia were grown with and without melanin inhibitors (30 µg/ml). Conidia exhibiting a brown phenotype were phagocytosed more readily by BMDM cells than those grown without melanin inhibitors (**Figure 2D**). After phagocytosis, BMDM were lysed, and the recovered intracellular fungal cells were plated on PDA medium for 3 d. Conidia grown in the presence of all tested melanin inhibitors showed increased susceptibility to macrophage fungicidal mechanisms (**Figure 2E**), indicating that inhibition of melanization led to increased *L. prolificans* susceptibility to macrophage killing.

#### Zeta Potential analysis

Melanins are known to be negatively charged (40, 41). Consequently, the zeta potential of the *L. prolificans* cells grown under different conditions could be used to evaluate whether the inhibitors were able to decrease melanization by assessing the resulting charge. The most negative zeta potential was observed for melanin ghosts from cells grown in PD only (no melanin inhibitors added) (**Table 1**). Cells grown with 30 µg/ml of pyroquilon and tricyclazole, both DHN-melanin inhibitors, displayed a brown cell phenotype and revealed increased zeta potentials approaching positive values (**Table 1**). In contrast, cells treated with niacin, a DOPA-melanin inhibitor, retaining a black cell phenotype, did not exhibit this change (**Table 1**). Thus, melanization inhibition was found to be accompanied by an increase in zeta potential.

**Table 1.** Zeta potential of melanin ghosts from *L. prolificans* cells grown in different conditions. Melanin inhibitors [30 µg/ml].

| Condition | Zeta Potential (mV) |
| --- | --- |
| PDB | (-25.00) |
| PDB + PYROQUILON | 0.62 |
| PDB + TRICYCLAZOLE | 3.11 |
| PDB + NIACIN | (-17.65) |

### Structural Characterization of *L. prolificans* Melanins

#### TEM Assessment of Melanin Deposition on the Cell Wall

TEM images from conidia grown in PDA media showed changes in deposition of the melanin layers on the *L. prolificans* conidia cell wall upon growth with 30 *µ*g/ml of each of the melanin inhibitors. In the presence of pyroquilon, tricyclazole, and the combination of pyroquilon and niacin, the melanin layers were reduced on the fungal surface (**Figure 3, arrowheads**), and this change was correlated with the brownish conidia phenotype, which differed from the darker/black phenotype observed for the native melanin (control) and with a niacin additive (**Figures 2A and 3**).

**Figure 3.**
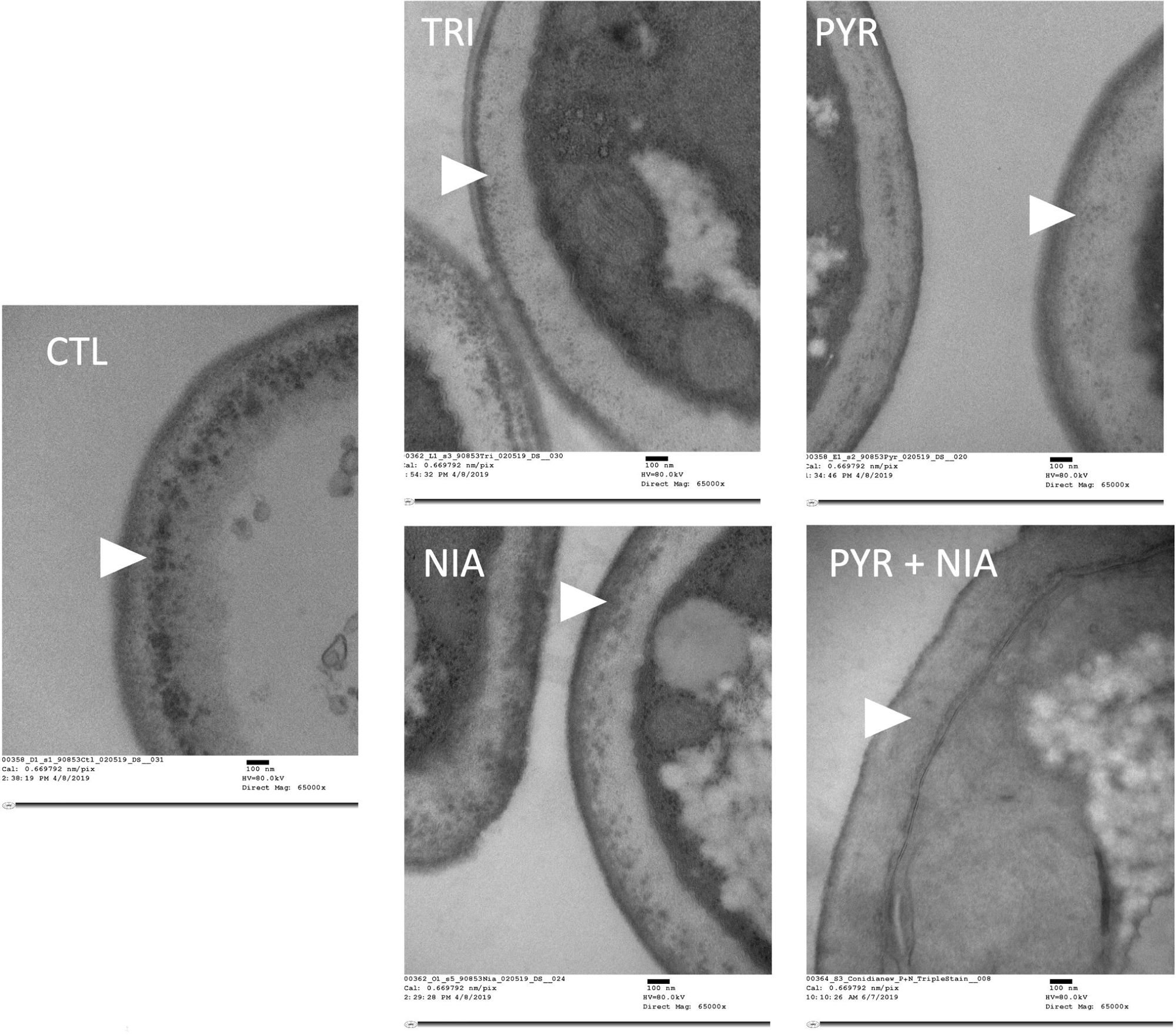
Structural analysis by TEM. Melanized cell wall of *L. prolificans* in the presence of 30 µg/ml of melanin inhibitors reveals variations in the deposition of melanin pigments. Representative cross-sectional transmission electron micrographs of *L. prolificans* show a close-up view of the cell wall, making it possible to see differences in layered melanin pigment arrangements among the conditions tested (white arrowheads). Melanin inhibitors. Scale bar, 100 nm. Control (CTL – no melanin inhibitor), pyroquilon (PYR), tricyclazole (TRI), pyroquilon + niacin (PYR+NIA), and niacin (NIA).

#### EPR Measurement of Free Radical Content

Melanins contain stable free radicals (31,32). EPR spectra of *L. prolificans* melanin in native form and after treatment with melanin inhibitors revealed a single slightly asymmetric line shape characteristic of DOPA-melanin (eumelanin) (21) (**Figure 4A**). Whereas the signal intensity was lower than native melanin upon treatment with pyroquilon and tricyclazole, the samples isolated after treatment with niacin displayed about sevenfold higher signal intensity (33) (**Figure 4A**).

**Figure 4.**
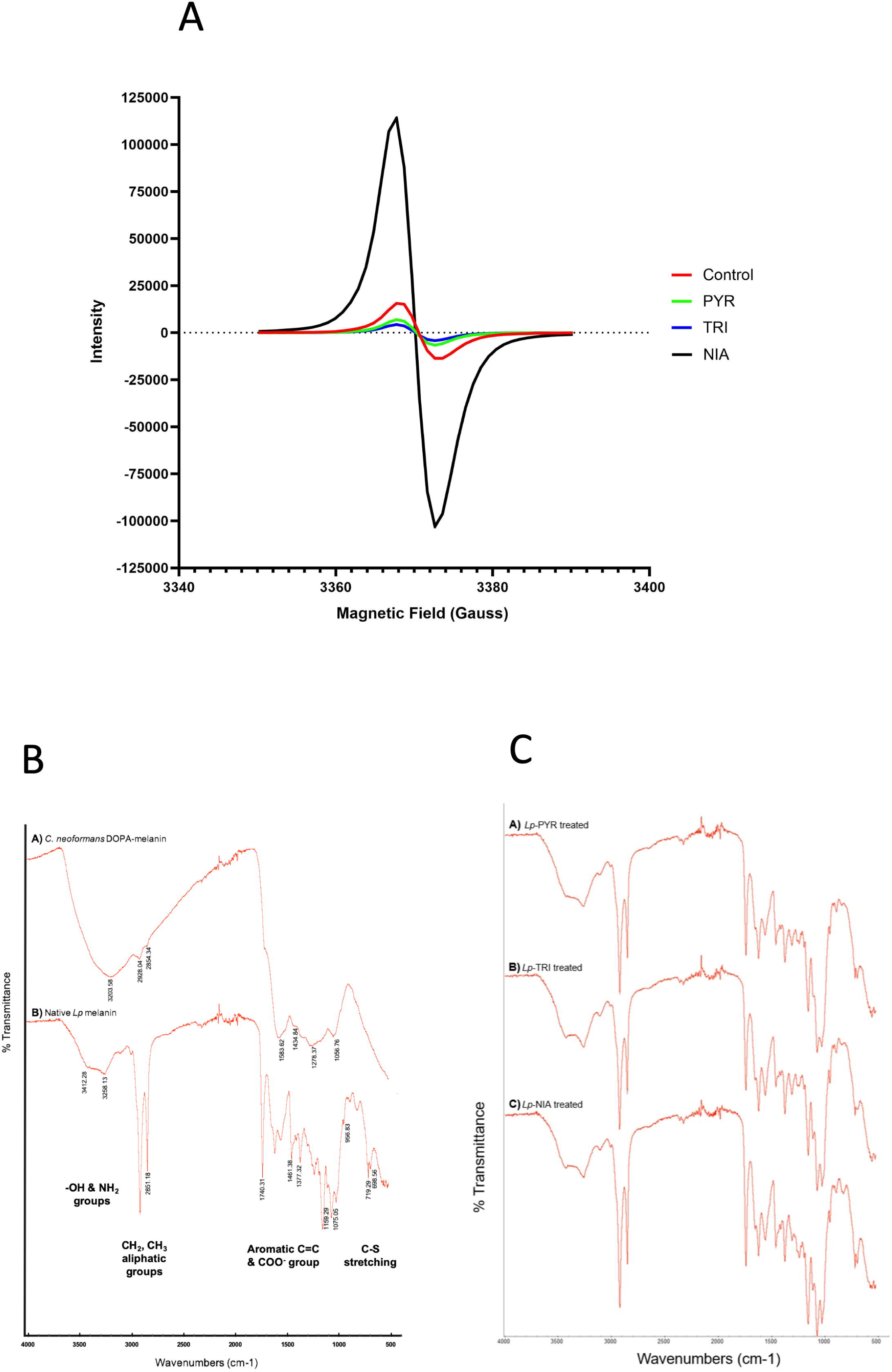
Structural analysis by EPR and FITR. Electron paramagnetic resonance (X-band, 9.43 GHz) spectra of *L. prolificans* melanins **(A)**. Pigment samples were isolated from fungal cultures grown with 30 µg/ml of the melanin inhibitors pyroquilon (PYR), tricyclazole (TRI), and niacin (NIA). The control sample corresponds to the native pigment synthesized without added inhibitors. Representative data from two independent experiments. Fourier Transformation Infrared Spectra (FTIR) spectra of *C. neoformans* DOPA-melanin and *L. prolificans* native melanin **(B)** and FTIR spectra of *L. prolificans* melanin isolated from fungal cultures grown with 30 µg/ml each of the melanin inhibitors pyroquilon (PYR), tricyclazole (TRI), or niacin (NIA), **(C)**. Representative spectra from two independent experiments are shown. CTL: control.

#### Identification of Chemical Groups by FTIR spectroscopy

The FTIR spectrum of native *L. prolificans* melanized cell walls (ghosts) displayed bands similar to *C. neoformans* DOPA-melanin (**Figure 4B**). A broad band centered around 3300 cm^-1^, indicating the presence of hydroxyl (-OH) and amine (NH2) groups (34,35), was observed from both samples. Features at 2950-2850 cm^-1^ corresponding to aliphatic CH2 and CH3 groups (35,36) were visible with higher intensity in the native *L. prolificans* melanin, as well as bands from aromatic C=C and COO^-^ groups in the region 1700-1380 cm^-1^ (37). The aliphatic and carbonyl features may be attributed to the cell walls to which the pigment is attached. The presence of bands around 700 cm^-1^ uniquely in the *L. prolificans* sample is diagnostic for pheomelanin (38). Melanin ghosts obtained from *L. prolificans* grown in the presence of the various inhibitors demonstrated the same spectral pattern as native melanin, indicating that the above-mentioned moieties remain present (**Figure 4C**). Thus, the IR profile of *L. prolificans* native melanin suggested a mixture of eumelanin and other pigments.

#### Elemental Analysis of *L. prolificans* Melanin Monomeric Units

5,6-dihydroxyindole (DHI) and 5,6-dihydroxyindole-2-carboxylic acid (DHICA) are established monomers of melanins (31). Depending on whether they are formed by the L-DOPA or other biosynthetic pathways, fungal melanin structures include either both DHI and DHICA monomers including of 6–9% nitrogen, or 1,8-dihydroxynaphthalene (DHN) with essentially no nitrogen (< 0.1%). For instance, elemental analysis of *Auricularia auricula* melanin revealed 41% C, 5% H, and 1.6% N, without significant S, a composition characteristic of pheomelanins, suggesting that it contained a mixture of DHICA/DHI and DHN eumelanin (22). Similarly, *L. prolificans* melanin from cells grown in PD medium contained 68% C, 9% H and 1.6% N, with no significant S, also suggesting a mixture of DHICA/DHI- and DHN-eumelanin (22) (**Table 2**). To gain insights into the biosynthetic pathways involved in the formation of melanin in *L. prolificans*, we measured the elemental composition of the pigments produced when the cultures were grown in the presence of various melanin synthesis inhibitors. The addition of 30 µg/ml of most melanin inhibitors did not change the melanin elemental composition significantly. However, the addition of pyroquilon and niacin together resulted in striking increases for N, O, and S, decreases in C and H, and a consequent shakeup in the C:N:O ratio (**Table 2**).

**Table 2.** Elemental Analysis of melanin ghosts from *L. prolificans* cells grown in PD with melanin inhibitors [30 µg/ml]. PYR+NIA: pyroquilon + niacin.

| Element | CONTROL | PYROQUILON | TRICYCLAZOLE | NIACIN | (PYR+NIA) |
| --- | --- | --- | --- | --- | --- |
| C | 68.08% | 67.06% | 65.65% | 69.68% | 48.25% |
| H | 9.55% | 9.57% | 9.60% | 9.18% | 7.40% |
| N | 1.63% | 1.58% | 1.93% | 1.70% | 4.31% |
| O | 19.45% | 20.40% | 21.71% | 18.87% | 39.02% |
| S | <0.5% | 1.40% | 0.87% | <0.5% | 1.16% |
| C : N : O | 42 : 1 : 12 | 42 : 1 : 13 | 34 : 1 : 11 | 41 : 1 : 11 | 11 : 1 : 9 |

#### ^13^C ssNMR Analysis of Melanized Cell Walls

To interpret the *L. prolificans* melanin NMR spectra, we first compared the control sample (without inhibitor) to synthetic DHN-melanin and its 1,8-DHN precursor. As expected from prior studies of melanization in *C. neoformans* (6,39), the CPMAS spectra of *L. prolificans* melanin ghosts included prominent resonances from retained cell-wall polysaccharides (50-110 ppm) and trapped neutral lipids (10-40 ppm). For the melanin pigments of primary interest in the current study, the resulting ^13^C NMR spectra displayed a broad pattern of resonances (including one relatively sharp feature at 130 ppm) in the 110-150 ppm aromatic spectral region, suggesting that *L. prolificans* ghosts contained either or both of the DHN- and L-DOPA-derived pigments (**Figure 5A**) (39). ^13^C ssNMR did not distinguish between these two melanin types since they have relatively similar spectral features. Nevertheless, the comparison was informative because the hydroxylated aromatic carbon resonances (140-150 ppm) appearing in spectra of synthetic DHN melanin and L-DOPA melanin (39) are absent in *L. prolificans* melanin.

**Figure 5.**
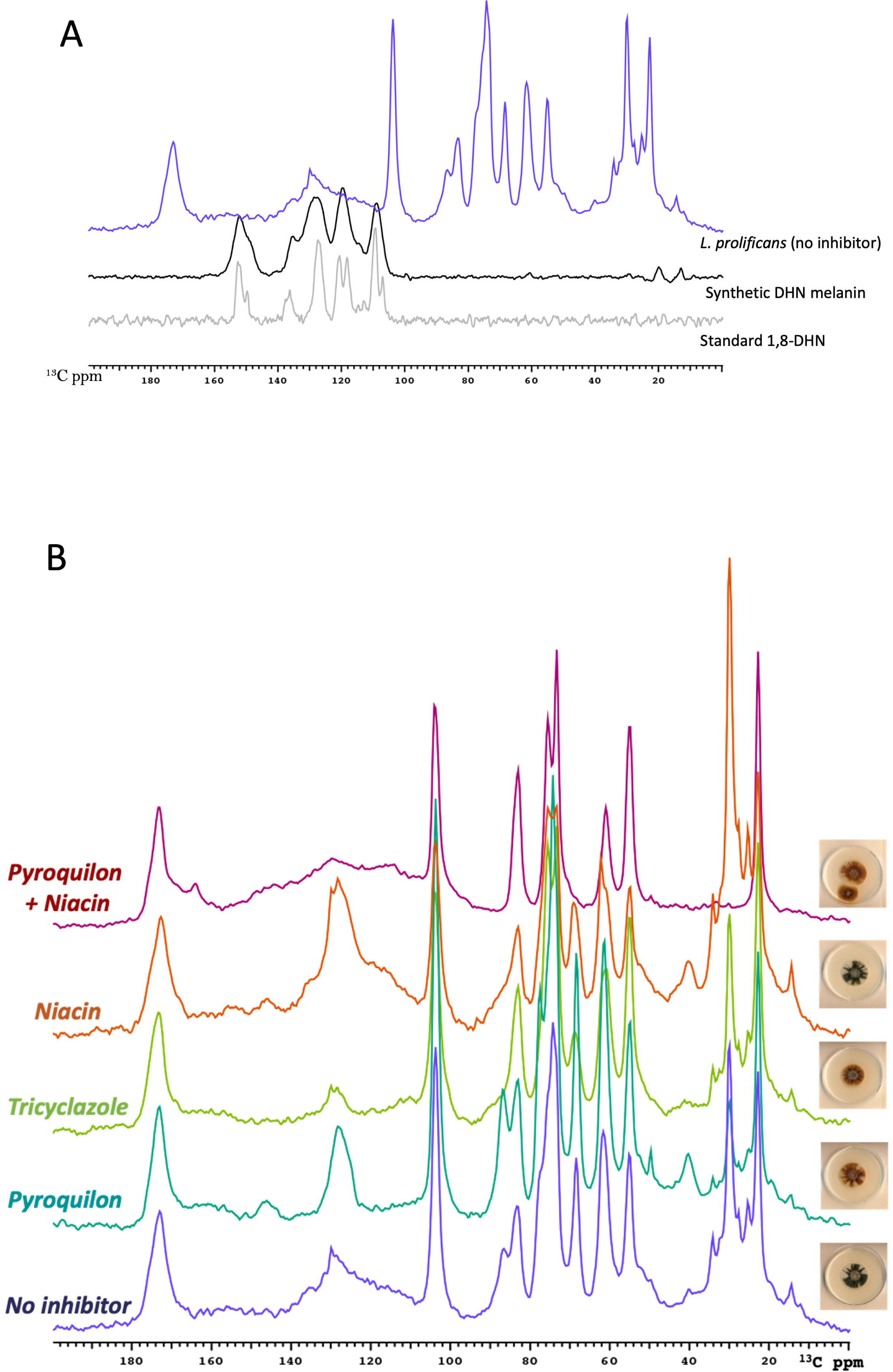
Structural analysis of ssNMR spectra. 150 MHz solid-state CPMAS ^13^C NMR spectra of synthetic DHN-melanin and its standard precursor compared with *L. prolificans* melanin ghosts **(A).** CPMAS ^13^C NMR spectra of melanin ghosts obtained from *L. prolificans* in the absence or presence of 30 µg/ml of each melanin inhibitor (**B**).

Turning the focus to the impact of inhibitors on the broad aromatic resonances between 110 and 150 ppm that are characteristic of amorphous melanin pigments, the ^13^C CPMAS NMR spectra *L. prolificans* melanin ghosts showed a broad group of aromatic NMR signals for the melanins cultured without inhibitors, but a narrower resonance for fungal ghosts cultured in the presence of pyroquilon or tricyclazole. By contrast, the addition of niacin produced narrow aromatic resonances that were visible above the broad spectral feature, and the pyroquilon-niacin combination yielded only the broad aromatic NMR signals similar to the control *L. prolificans* melanin (**Figure 5B**). The key regions of the ssNMR spectra in **Figure 5B** and a comparison between the characteristics and composition of *L. prolificans* melanin without or with 30 µg/ml of each melanin inhibitor are summarized in **Table 3**. Notably, the persistence of defined melanin-like aromatic NMR features upon addition of a single inhibitor, as also found in the EPR data of **Figure 4A**, could be attributed to a compensation effect, whereby the fungus accesses alternative precursors and pathways when a particular pathway is blocked.

**Table 3.** Summary of ^13^C CP data of melanin ghosts from *L. prolificans* cells grown in different conditions. Melanin inhibitors [30 µg/ml].

| Sample (inhibitor) | Carbonyl | Aromatic Broad | Aromatic Narrow | 88 ppm | 70 ppm | 50 ppm | 40 ppm | Aliphatic |
| --- | --- | --- | --- | --- | --- | --- | --- | --- |
| No Inhibitor | 173 ppm | Broad 100-145 ppm | Short 30 ppm | Present | Present | Very small | Very small intensity | High intensity 30 ppm |
| Pyroquilon | 173 ppm | Smallest area of all samples; shoulder at 148 ppm | More intense than control | Present at higher intensity than control | Present; higher intensity | Higher intensity than in control | Highest intensity from all samples | Very low intensity |
| Tricyclazole | 174 ppm<br>Slightly more intense than in control | Very small compared to control; no 148 ppm shoulder | Less intense than control | Becomes a shoulder | Present | Not present | Kind of there | Lower intensity than control |
| Niacin | 172 ppm | Greater area than control | More intense than control | Becomes a shoulder; | Present | Not present | Present | Higher intensity than control |
| Pyroquilon + Niacin | 172 ppm, broad extends to 158 ppm; 2<sup>nd</sup> peak at 163 ppm | Broader region and greater area from all samples | Less intense compared to control | Completely gone | Completely gone | Almost completely gone | Not present | Not present |

#### Optical and ssNMR Assessment of Pyomelanin as a Possible *L. prolificans* Melanin

To obtain a better understanding of melanin production in *L. prolificans*, we grew cells in the presence of the accepted substrate for pyomelanin, homogentisic acid (HGA), and its nitisinone inhibitor (20). Alkaline solutions of the resulting melanins exhibited a strong optical absorbance in the UV region, which gradually decreased toward longer wavelengths (range of 400-700 nm). Alkali-soluble pigments from cell cultures grown in the presence of different melanin inhibitors and HGA indicated that pigment formation was increased by addition of 1 mM HGA but decreased by 30 µg/ml of nitisinone (**Figure 6A**). We then used ^13^C ssNMR to analyze *L. prolificans* melanin ghosts grown with 30 µg/ml each of three different inhibitors designed to block the three known fungal melanin pathways. Melanin production was suppressed when *L. prolificans* cells were grown in the presence of pyroquilon (DHN-melanin pathway inhibitor), niacin (DOPA-melanin pathway inhibitor) and nitisinone (HGA-melanin pathway inhibitor), as evidenced by diminished ^13^C NMR spectral contributions in the 110-155 ppm range compared to control (no inhibitors). Additionally, neutral lipids (30 ppm) were diminished, leaving the retained polysaccharides as the most abundant constituents (50-110 ppm) (**Figure 6B**).

**Figure 6.**
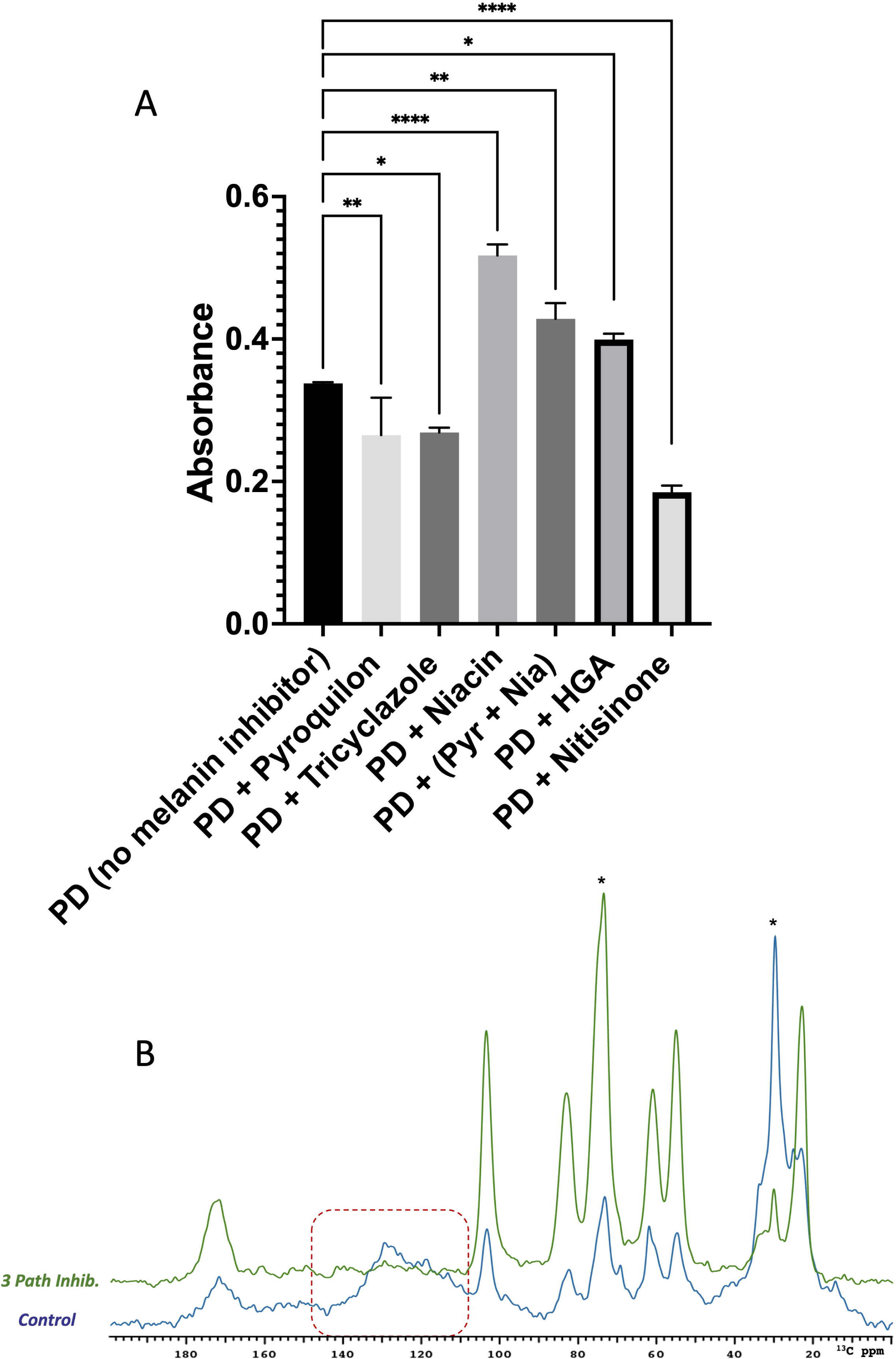
Melanin production upon addition of melanin inhibitors and homogentisic acid. Melanin absorption spectra acquired in the absence or presence of melanin inhibitors **(A)**. These experiments were repeated three times in duplicate and standard deviations are calculated from three independent experiments. *L. prolificans* was grown in PD liquid media without or with 1 mM of the HGA melanin substrate and in the absence or presence of 30 µg/ml each of one or more melanin inhibitors. Quantification of the pigment was measured from its absorbance at 405 nm (**A**). This experiment was repeated three times in duplicate and standard deviations are calculated from three independent experiments. ^13^C CPMAS ssNMR spectra of *L. prolificans* with no inhibitors (blue) or supplemented with 3 pathway inhibitors (green: DHN, DOPA and HGA) **(B)**. Pyr + Nia: Pyroquilon + Niacin. HGA: homogentisic acid. * *p* < 0.05, ** *p* < 0.005, **** *p* < 0.0001.

## Discussion

Although invasive *L. prolificans* infections remain relatively rare at present, they are high consequence events due to their association with high mortality rates and this species’ significantly strong resistance to antifungal drugs. The combination of high mortality, high antifungal resistance, and disease in immunocompromised hosts poses a tremendous therapeutic challenge. The basis for this antifungal drug resistance is unknown, but attention has focused on melanin, as it reduces the susceptibility of other fungal species to drugs such as amphotericin B (40). Nonetheless, a study of the potential role of melanin conducted by deleting genes involved in DHN melanin production found no effect on amphotericin susceptibility, even though increased vulnerability to oxidative killing was observed (14). Given that some fungi are known to make more than one type of melanin, we were prompted to investigate the types of melanin produced by *L. prolificans*. Considering the analytical challenges of characterizing these insoluble amorphous biopolymer composites structurally, we sought to assemble a suite of complementary information using indirect approaches such as the use of inhibitors, the analysis of degradation products, and the tools of molecular biophysics.

Some species of *Scedosporium*/*Lomentospora*, such as *Scedosporium* (formerly *Pseudallescheria*) *boydii* (17) and *Scedosporium apiospermum* (41), produce melanin via the DHN pathway, yielding a type of melanin also found in *A. fumigatus* conidia; however, the characterization and functions of melanins from these species have been sparsely studied to date (7,14,17). Previous reports showed that in *S. boydii*, a species related to *L. prolificans,* a clear difference could be seen in the color of conidia isolated from cultures grown in the presence of DHN-melanin inhibitors (17). Furthermore, a UV/visible spectral analysis of melanin extracts confirmed that DHN-melanin is a major pathway to produce melanin in *S. boydii* (17). Notably, Thornton and collaborators developed the mAb CA4, where its antigen was identified as the melanin biosynthetic enzyme 4HNR (from the DHN pathway) (16). Their data showed that enzyme-deficient mutants produced orange-brown or green-brown conidia, in contrast to the black wild type conidia, but the mutant conidia were not albino/colorless (16). In addition to DHN-melanins, pyomelanin and DOPA-melanin are synthesized by *A. fumigatus* (7,42), which deposits melanin on the cell surface, similar to *L. prolificans.* We hypothesized that the melanin pathways could be similar between these species.

In the current work, we characterized *L. prolificans* melanin chemically, biophysically, biologically, and structurally in the context of possible inhibition of particular pathways for the biosynthesis of DHN- DOPA- and pyomelanin. For *L. prolificans,* the color/pigmentation of conidia cells were each influenced by the choice of media and the age of the culture. The addition of three melanin inhibitors together (pyroquilon, niacin and nitisinone) decreased the UV absorbance of alkaline extracts in a manner related to melanin discoloration. High doses of pyroquilon (DHN-melanin inhibitor) and niacin (DOPA-melanin inhibitor) significantly decreased the pigmentation of the *L. prolificans* cells and the thickness of melanin layers observed on the fungal surface by TEM. Conversely, the addition of the pyomelanin substrate HGA increased pigment formation.

Elemental analysis supported a mixture of DHICA/DHI and DHN eumelanin (17). No sulfur was found in most culture conditions, so most of the pigments could be classified as eumelanins. Nonetheless, the addition of both DHN- and DOPA-melanin inhibitors produced a melanin that was rich in nitrogen and oxygen but deficient in carbon. Laccase enzymatic activity was evident for *L. prolificans,* indicating that this species can produce more than just DHN-melanin, marking the first report of it. Since melanin is a negatively charged polymer, the zeta potential measurements provided a means to determine whether various substrates were equally incorporated into melanin. For melanins synthesized in the presence of both DHN-melanin inhibitors, which resulted in brown cell phenotypes, the usual negative zeta potential shifted to positive values, suggesting significant alteration of the melanin pigment structure. Moreover, the melanin-linked anti-phagocytic role observed in several fungal infections (7) was reduced for the brown conidia produced upon the addition of DHN and DOPA melanin inhibitors, making these fungal cells more susceptible to killing by BMDM cells. Complementing these biological and biophysical measurements, structural analyses using EPR and FTIR also suggested that *L. prolificans* melanin is a mixture of eumelanin and pheomelanin. Finally, ssNMR demonstrated the suppression of production for melanin as well as associated lipids in melanized cell walls only when two or three melanin inhibitors were added in concert. This responsiveness to a cocktail of multiple inhibitors also supports the notion that *L. prolificans* can utilize several melanization pathways, offering compensatory protection if a single path is inhibited; pigment formation would then be suppressed only if all biosynthetic options are blocked.

In summary, the current work offers new insights into melanin production in the human pathogenic fungus *L. prolificans*. Many fungal species are known to produce different types of melanins (7), and *L. prolificans* appears to belong to this group by virtue of its ability to access the different pathways for producing DHN-, DOPA- and pyomelanin pigments. That said, our inability to completely inhibit melanization with known inhibitors of these pathways, evidenced by distinctive attributes observed spectroscopically, raises the possibility that *L. prolificans* makes yet another type of melanin, which can co-exist with a mixture of the DHN-, DOPA- and pyomelanin types.

## Data Availability

All data is contained within the manuscript. Data is available to share upon request. Please contact the corresponding author.

## Supporting Information

This article contains no supporting information.

## Acknowledgements

We are thankful for the support of the Analytical Imaging Facility at The Albert Einstein College of Medicine. The authors are grateful to their colleagues, Dr. M. Wear for helpful discussions, and Van Chanh Phan and Yuliana Dominguez Paez for helpful experiments of NMR.

## Author contributions

Conceptualization: LLL; Investigation: LLL, EC, CC, AK; Visualization: LLL, CC, EC; Supervision: AC, RES; Writing–original draft: LLL; Funding acquisition: AC, RES. All authors have read and approved the final manuscript.

